# Route Learning and Transport of Resources during Colony Relocation in Australian Desert Ants

**DOI:** 10.1101/2024.08.27.609847

**Authors:** Sudhakar Deeti, Donald James McLean, Trevor Murray, Ken Cheng

## Abstract

Many ant species are able to respond to dramatic changes in local conditions by relocating the entire colony to a new location. While we know that careful learning walks enables the homing behavior of foraging ants to their original nest, we do not know whether additional learning is required to navigate to the new nest location. To answer this question, we investigated the nest relocation behavior of a colony of Australian desert ants *Melophorus bagoti* that relocated their nest in response to heavy rainfall in the semi-desert terrain of Alice Springs. We identified five types of behavior: exploration between nests (Old-to-New nest and New-to-Old nest), transport from Old to New nest, and re-learning walks at Old and New nests. Initially, the workers performed relearning walks at the Old nest and exploratory walks between the Old and New nests. Once they completed the exploratory walks, the workers transported resources and brood to the new nest. Finally, we observed the workers performing relearning walks at the New nest. While the relearning walks at the Old nest were slow and appear to enable exploratory walks to the New nest, the relearning walks at the new nest were faster and appeared to enable homing from foraging trips. These observations shed insight on how learning helps these ants to respond to sudden changes in their environment.

**Statement of significance:** In this study, we examined one nest of red honey ants, *Melophorus bagoti*, in Central Australia relocating their nest after a spell of heavy rainfall, which, we presumed, damaged the nest that the colony was living in. After the new nest some 20 m away was mostly dug, we examined the behaviors that orchestrated the move. Workers did a bit of learning of the route to the new nest before transporting any resources. They executed short loops around the old nest, mostly aimed in the direction of the new nest, known as relearning walks. Then they walked back and forth between the old and new nests without carrying any resources, a process we called exploration walks. Then the workers undertook the laborious process of moving house, that is, transporting workers unfamiliar with the route, larvae, and pupae to the new nest. After this stage, we observed ants doing more relearning walks at the new nest in preparation for foraging from their new home. These relearning walks covered all directions around the nest. Our study showed for the first time the importance of the process of learning in nest relocation, a key response to extreme environmental challenges, broadening the role of learning in the lives of these social insects.

## Introduction

For social insects like ants, social wasps, and honeybees, nests are a vital resource. They serve as a sanctuary, offering protection, storage space, and a communal platform (Jeanne, 1975; Seeley & Morse, 1976; Tschinkel, 2004; Wilson, 2000). Sometimes, however, nests must be abandoned and the colony must relocate due to factors such as disturbances, microclimate changes, predation, and competition (Hölldobler & Wilson, 1990; McGlynn, 2012). Transport of workers and brood toward new nests is not restricted to colony relocation; it also occurs in the context of the foundation of new colonies by colony fission, as seen in species like *Cataglyphis cursor* (Chéron et al., 2011). Nest relocation in desert ants is a remarkable behavior that encompasses collective action, navigation, and adaptation to the extreme conditions of the desert environment. While this study focuses on the red honey ant *Melophorus bagoti*, the desert ant *Cataglyphis iberica* provides a compelling example of sophisticated navigational strategies and diverse nest relocation tactics. This species exhibits exceptional navigational abilities, employing a combination of visual cues—such as the position of the sun—and path integration, which involves tracking their movements relative to their starting point (Fourcassié et al., 2000). These ants maintain remarkable accuracy in their navigation despite the harsh desert terrain, effectively integrating visual information with internal movement tracking to relocate their nests. The diverse strategies for nest relocation observed in *Cataglyphis iberica* include following prominent landmarks and adjusting their navigation based on environmental changes (Fourcassié et al., 2000).To date, only four sets of field-based studies have investigated nest relocation in desert-living ants: on *Cataglyphis iberica* in Spain (Dahbi et al., 2008), on *Melophorus bagoti* (Deeti & Cheng, 2021a; Schultheiss et al., 2010), and on nocturnal bull ants *Myrmecia Midas* (Deeti et al., 2024a). While these studies documented the relocation process, which includes the physical movement of the nest and the associated logistical coordination, they did not explore the learning processes involved in navigation to the new nest site. Specifically, they did not address whether ants engage in exploratory behavior or ‘learning walks’ that facilitate successful nest relocation. Understanding these strategies is crucial as it offers insights into how ants manage complex navigational tasks and adapt to extreme conditions, which can be valuable for studying similar behaviors in other species and environments.

Most ant species rely heavily on trial pheromones and not or very little on visual cues. Hymenopterans, such as bees, wasps, and some ants, learn and relearn the visual cues surrounding their nest for navigation (Collett & Zeil, 1996; Lehrer & Bianco, 2000; Nicholson et al., 1999; Philippides et al., 2013; Zeil et al., 1996). These learning walks and flights enable these insects to efficiently navigate in their environment. The learning or orientation flights of bees and wasps play a key role in learning about the location of landmarks, food sources, and potential hazards (Collett & Lehrer, 1993; Zeil, 1993). Individual naive ants explore their environment in a systematic pattern in order to learn the location of their nest before beginning their foraging life (Deeti & Cheng, 2021b; Fleischmann et al., 2016; Jayatilaka et al., 2018). Desert ants also perform learning walks at feeders and also learn the route (Nicholson et al., 1999; Vermehren et al., 2020). Experienced ant foragers also sometimes perform short looping walks reminiscent of learning walks, called re-learning walks (Jayatilaka et al., 2018; Narendra & Ramirez-Esquivel, 2017), for example, at the start of a new day or when presented with a new landmark set up (Wystrach et al., 2014). Relearning flights or walks are also important for navigation in wasps, bees and ants. During relearning flights, the insect typically flies in a looping or circuitous pattern, similar to learning walks seen in ants (Collett & Zeil, 1996; Lehrer & Bianco, 2000; Nicholson et al., 1999; Philippides et al., 2013; Zeil et al., 1996), where insects update their memories of their surroundings seemingly to adjust to changes in their environment or degradation of their memories. Updating is important: studies on two species of bull ants of *Myrmecia* genus showed that changes in the landmark panorama affected the visual navigation ability of these nocturnal ants. These changes prompted ants to repeatedly turn back and look toward the nest while heading out to forage (Islam et al., 2020; Narendra & Ramirez-Esquivel, 2017). While we know much about the naïve learning walks, we know less about when and how foragers augment their initial learning as they experience navigational challenges, changes to their visual environment, or locations beyond their familiar range, such as during nest relocation.

Relocation in social insects differs from relocation in solitary animals. While both require searching for a new nest, social insects must also coordinate the transfer of colony members unable to make the trip alone. This intricate navigational behavior has been extensively studied in honeybees (Seeley, 2009) as well as in a few species of wasps (Strassmann, 2001) and ants (Franks et al., 2005). During nest relocation in colonies with monomorphic workers, different workers take on different roles and perform specific tasks to ensure a successful move (Visscher, 2007). Scouts explore the surroundings to find potential nest sites, assess these sites, and communicate their findings to the colony (Pratt et al., 2005). Excavators or builders construct new chambers at the chosen location (Beshers & Fewell, 2001). Transporters coordinate the transportation of brood, including eggs, larvae, and pupae, to the new location (Visscher, 2007). It remains unclear, however, whether navigational learning is required to support these behaviors and enable these individual tasks, which often require travel between the old and new nest locations.

Here, in a natural experiment, we observed the relocation behavior of one rain-damaged colony of desert ants. We filmed the behavior of these ants around their old nest, their new nest, and during their trips between the two nests to examine the path structures they follow and whether they transport brood during nest relocation. Beyond describing these behaviors, we formed the following research questions before analysing any of the data: Do colony members perform relearning walks before engaging in relocation behaviors? Do colony members perform any additional learning walks at the new nest, once they have travelled there? What is the structure of these learning behaviors and to what extent do they differ from naïve learning walks? Answering these questions will help us to better understand the role that learning plays in relocation, while also shedding light on how life-long learning can help facilitate navigational behavior across the wide range of environments and scenarios that these animals find themselves in.

## Materials and Methods

### Study species

All observations were performed in an open and grassy arid desert land at the Centre for Appropriate Technology in Alice Springs from November to December 2022. The region is characterized by vast semi-desert landscapes with sparse vegetation, such as *Spinifex* grass, *Acacia* trees, shrubs, and *Eucalyptus* trees. The red honey ant, *M. bagoti*, is the most thermophilic ant species found on the Australian continent (Christian & Morton, 1992). Our observations revealed that these ants were usually active outside the nest when temperatures ranged between 32 and 40°C. During the hot southern summer, these *M. bagoti* ants engage in foraging. They scavenge primarily dead spiders, sugar ants, termites, centipedes, moths and other dead arthropods. Additionally, they also collect sugary plant exudates and seeds (Muser et al., 2005; Schultheiss & Nooten, 2013). These Australian desert ant *Melophorus bagoti* colonies are widely distributed across the central Australian semi-desert, where these endemic honey ants make their monodomous colonies. The ants are known for their unique nesting habits, which involve excavating subterranean nests with multiple chambers made in sandy soil with typically only one entrance on the ground (Deeti et al., 2023; Deeti et al., 2020). The red soil was composed of sand sediments laid down by floods (Baker et al., 1983). During summer months, we had the opportunity to record the workers’ activity in a collapsed colony after a heavy rainfall. We found the workers of the colony excavating the new nest a week after heavy rain. First seven days foragers were found excavating a new tunnel at the Old nest. Later on, we found the foragers were scouting and excavating the new nest. We have started out structural recordings and relocation related events of the nest from the 8^th^ day we started to record the relearning walks of the worker ants at the Old nest. While Australian ethical clearance is not required to conduct research on the ants, our study was completely observational and did not impose any manipulations during the nest relocation.

### Procedure

The methods for this study involved observation and documentation the structure of the paths of workers during nest relocation. The study was initiated after the discovery of the workers excavating a new nest 20 meters away from the old nest three days after the heavy rainfall. After the discovery, we focused on key behaviors and events that were relevant to nest relocation, such as worker interactions, the process of nest relocation, worker roles and responsibilities, and any notable changes in behavior following heavy rainfall. In order to accurately document the behavior of workers during nest relocation, the study employed a systematic approach to observe and record events. The observed ant behaviors were classified into five different behaviors: Relearning (Old nest), Relearning (New nest), exploratory walk (Old-to-New), exploratory walk (New-to-Old), and transportation walks (always Old to New nest). Furthermore, these five behaviors have been grouped into two main categories: Nest-based learning walks and travelling-based learning behavior. In the Traveling based behavior category, ants started on a journey from one nest and ended it at the other nest. In contrast, during the nest-based learning behavior, ants initiated their journey at one nest and ended it at the same nest. This categorization allowed for a clear analysis of the ants’ behaviors in different scenarios during the nest relocation process. Nest based learning walks were defined as systematic loops around the nest area, both at the old and new nest locations, starting and ending at the same nest. Ants engaged in relearning walks did not carry debris, and these walks extended within a maximum distance of up to 30–80 cm from the nest entrance. We also observed ants traveling toward the Old or New nest. Ants which were shuttling between the Old and New nests without carrying any ant colony resources were considered as doing exploratory walks. On the other hand, when a worker carried pupae, callows, or larvae from the Old nest to the New nest, we considered it a Transporter. We recorded the one relearning walk of each unique individual worker (Old nest: N = 10; and New nest: N = 10), by painting them once they completed a relearning walk and only analysing the walks of unpainted ants. Then, we recorded the individual workers’ Travelling-based learning behavior from the Old to the New nest (N =16) and from the New to the Old nest (N = 10) and later when the workers’ transporting of resources and brood from the Old to the New nest (N = 10) at a midpoint 10 meters from the Old and New nests. In the travelling-based learning behavior we only recorded a meter of their paths during their travel to target nest at the midpoint. Once recorded their walks captured at them at the target location and painted with a dot of color on their gaster to avoid rerecording of the same ant.

### Recording

A Sony (4K) Handy Cam recorded the relearning walks. We placed the camera on a tripod at the Old nest to capture the relearning walks occurring there. On the same day we placed a second camera at the midpoint approximately 10 meters away from both the Old and New nests. Here, we recorded the exploration behavior of the ants as they ventured between the nests in both directions. We recorded the ants that happened to travel under the camera, capturing their activities during this exploration. On the next day, we placed a camera at the midpoint to record foragers transporting resources to the New nest, at the same location where the exploration behavior was earlier captured. On the day after the relocation was completed, we used the second camera to record the relearning walks of ants at the New nest. This comprehensive recording approach allowed us to study the workers’ behavior during all the crucial stages of nest relocation. Each recording was captured at 25 frames per second, and the recording area covered a 1-square-meter area with a resolution of 3860*2160 pixels. To record the individual ant relearning behavior at the Old and New nests, we observed the relearning walks as the ants left the nest and continued observing them until they returned back to the nest. For exploratory walks, we followed the ants from the Old and New nests, recorded their behavior at the midpoint, and continued to follow them until they arrived at their goal. As for the transporters, we followed them from the Old nest until they reached the recording area, where we filmed their behavior, and continued to follow them until they entered the new nest. To avoid re-recording ants in all experimental observations, we painted each videotaped ant’s gaster with a color (Citadel) immediately before it entered its destination nest. These painted ants were found frequently exploring between the Old and New nests. However, we did not record any of their activities in these further appearances.

### Tracking

We used the animal tracking program DLTdv8 (8.2.9) Matlab (2022B) to extract frame-by-frame coordinates of the head and thorax of each ant in each video from our recordings of their behavior (S.Figure1). These frame-by-frame coordinates were used for all subsequent analyses of the workers’ trajectory structural behavior during nest relocation. The thorax positions were used for all the path characteristics described next and presented in Results while the head positions defined the direction in which an ant was facing (line from thorax through head), a variable that was not analysed in this study.

### Learning walks

For comparison, we recorded 10 red honey ants’ naïve, first learning walks at a different nest in the same area that did not relocate. Naïve ants were identified by painting every emerging ant from the nest with a dot of paint (Citadel brand) for nine days. After nine days, any unpainted ant emerging was considered naïve. A video camera was set up centred over the nest in the same fashion as at the Old nest. The camera was turned on as soon as an unpainted ant emerged. Characteristics of learning walks in this species, including the first, naïve walk, have been described (Deeti & Cheng, 2021b), but we obtained a new set here on statistical grounds: to avoid multiple comparisons with the same data set from Deeti and Cheng (2021b).

### Data Analysis

We analysed the ants’ trajectories (*N =* 56) using multiple indices of straightness and speed to understand the structure and function of these nest relocation behaviors. To understand how directed the ants’ trajectories were, we used three indices of straightness: *path straightness, sinuosity*, and 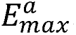, each of which relates to the directness of navigation toward a destination. We excluded the relearning walks in these analyses because they start and end at the same point. The simplest of these is *straightness*, calculated as the straight-line distance between the start and end of the path divided by the length of the path (Batschelet, 1981; Deeti et al., 2023; Islam et al., 2021; Islam et al., 2023; Lionetti et al., 2023). Straightness ranges from 0 to 1, with larger values indicating straighter paths, while smaller values indicate more curved or convoluted paths. *Sinuosity* is an estimate of the tortuosity in a path, calculated as 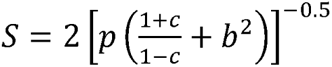, where *p* is the mean step length, *c* is the mean cosine of turning angles and *b* is the coefficient of variation of the step length. A trajectory *step* is the movement between successive recorded thorax positions, which is assumed to be a straight line. Accordingly, step length is the Euclidean distance between consecutive points along a path, and the turning angle refers to the change in direction between two consecutive steps. Sinuosity varies between 0 (straight) and 1 (extremely curved) (Benhamou, 2004). The maximum expected displacement of a path, 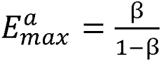, where β is the mean cosine of turning angles, is a dimensionless value expressed as a function of number of steps, and is consistent with the intuitive meaning of straightness (Cheung et al., 2007). Larger maximum expected displacement values indicate straighter paths, hence greater displacement, while a smaller value suggests more localized or constrained movement. Paths were characterized and visualized in R (version 4.2.1; R Core Team, 2020) using the packages trajr (McLean & Skowron Volponi, 2018) and Durga (Khan & McLean, 2023).

To understand how quickly and far the ants moved, we calculated speed, and three indices of movement in the relearning walks. Speed refers to the magnitude of an ant’s velocity and was calculated as the average over the entire trajectory for each ant, excluding the stopping durations (more than a frame). For movement characteristics, we calculated the maximum displacement, duration and area covered by the ants during the relearning walks. The nest location was chosen as the origin (0, 0). For each relearning walk, we noted the farthest distance the ant travelled from the nest entrance (maximum displacement). This helped us understand how extensively the ants explored their surroundings. We also recorded the exact duration of each relearning walk, from the moment the ant left the nest until it returned. For the area covered during these walks, we calculated the enclosed area, using the convex-hull method to encompass all the points visited by the ant. This area measurement provided insights into the spatial extent of their exploration.

### Statistical analysis

We used Generalized Linear Models (GLM) to understand the differences in path characteristics between the five nest relocation behaviors we observed. Relearning walks, however, are not comparable with travels between nests because the former start and end at the same place whereas the latter take the ant from one place to another. We thus split our models into two groups: Travelling and Relearning. Our travelling models compare the effect of walk types (New-to-Old nest, Old-to-New nest, and Transportation walks) on path characteristics. Relearning models, on the other hand, compare the relearning walks at the Old nest, at the New nest, and the first learning walks at the non-relocating nest. We used four models to compare the effect of the type of travelling behavior on path straightness, sinuosity, 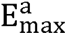 and speed. For Relearning walks we used four models which analysed speed, maximum distance, duration, and exploration area as our response variables. The GLM was formulated using the lme4 package (version 1.1-27) and fitted using the glmer function. Since path straightness and sinuosity are a bounded measures (0–1), we used the binomial family, whereas all the other response variables are only bounded by zero, and so for them we applied the Gaussian family of models. We employed the Tukey post hoc test (emmeans package) to perform pair-wise post hoc comparisons for each of our models. We report all significant terms and the significance values for all key pair-wise comparisons. Since we tested multiple dependent variables that are not totally independent of one another, we adopted p = 0.01 as an alpha level to lower type 1 errors. Statistical analyses were conducted using R (version 4.3.1). We visualized the data using box plots and drawings of paths.

For circular distributions, we tested their uniformity with the Rayleigh test (Batschelet, 1981). In case of significant non-uniformity, we also tested whether the distribution was significantly oriented in the goal direction using the V test. The V test returns a significant result when the distribution is significantly oriented in a theoretical (goal) direction. Circular statistics were conducted in R using circular package (Version 0.5-0).

## Results

In observing the behaviors associated with nest relocation, we identified a sequence of behaviors: firstly, Old-nest relearning walks, then Old-to-New and New-to-Old nest exploration, followed by Old-to-New nest transportation, then finally New-nest relearning walks. During the Old-nest relearning walks, ants leaving the nest travelled in the vicinity of the Old nest. After performing the relearning walks at the Old nest, workers moved back and forth between the Old and New nests. The ants then transported resources to the New nest. Subsequently, as the colony became established at the New nest, workers engaged in relearning walks at the New nest. We next detail characteristics of these behaviors.

### Exploratory and transportation walks

The analysis of ants’ walks between the two nests reveals significant differences in curvature in the paths of transportation walks and exploratory walks (Figures 1 to 3). All the walks, however, were headed in the general direction of the goal (Figure 1). The Rayleigh uniformity test and the distribution of headings with the *V* test showed significance in centroid headings (Figure 2) toward the goal in New-to-Old Exploration walks (Rayleigh: Z = 5.24, *p* ≤ 0.001; *V* test: Z = 2.9, *p* ≤ 0.008), in Old-to-New Exploration walks (Z = 9.6, *p* ≤ 0.001; *V* test: Z = 7.84, *p* ≤ 0.001) and in transportation walks (Z = 8.4, *p* ≤ 0.001; *V* test: Z = 6.72, *p* ≤ 0.001).

**Figure 1.**
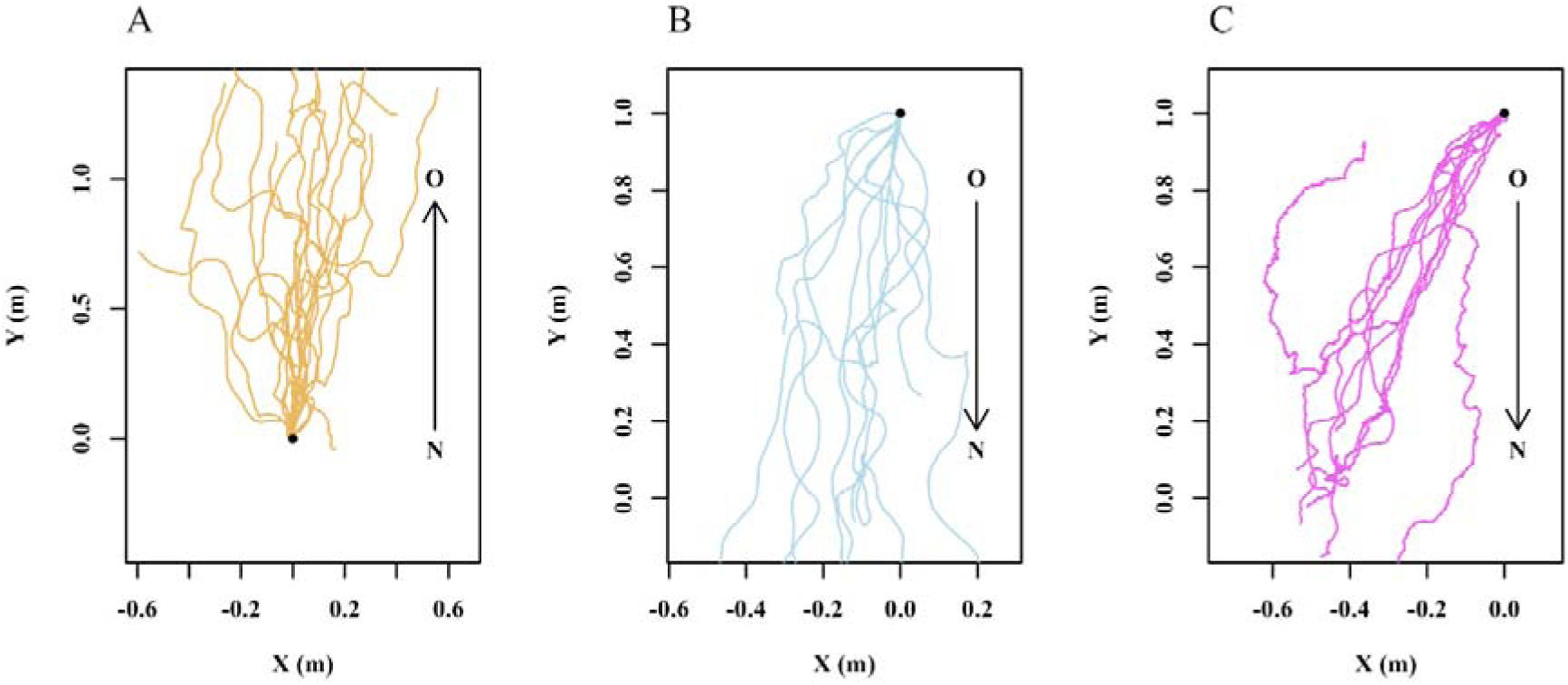
Trajectory plots of ants accomplishing different types of travel. (A) Ants returning to the Old nest from the New nest (N ≥ 10). (B) Ants heading to the New nest from the Old nest (N = 10). (C) Ants carrying brood and larvae to the New nest from the Old nest (N = 10). Each line represents the path of an individual ant. The trajectories are translated so that they all originate from position x = 0 and either y = 0 (first instance of the ant in the recording area) for ants heading to the Old nest, or y = 1 for ants heading to the New nest. The *x* and *y* coordinates represent the spatial position of the ants; spatial units are m. Arrows indicate expected direction of travel, from New (N) to Old (O) or from Old to New at 10 m away from both nests.

**Figure 2.**
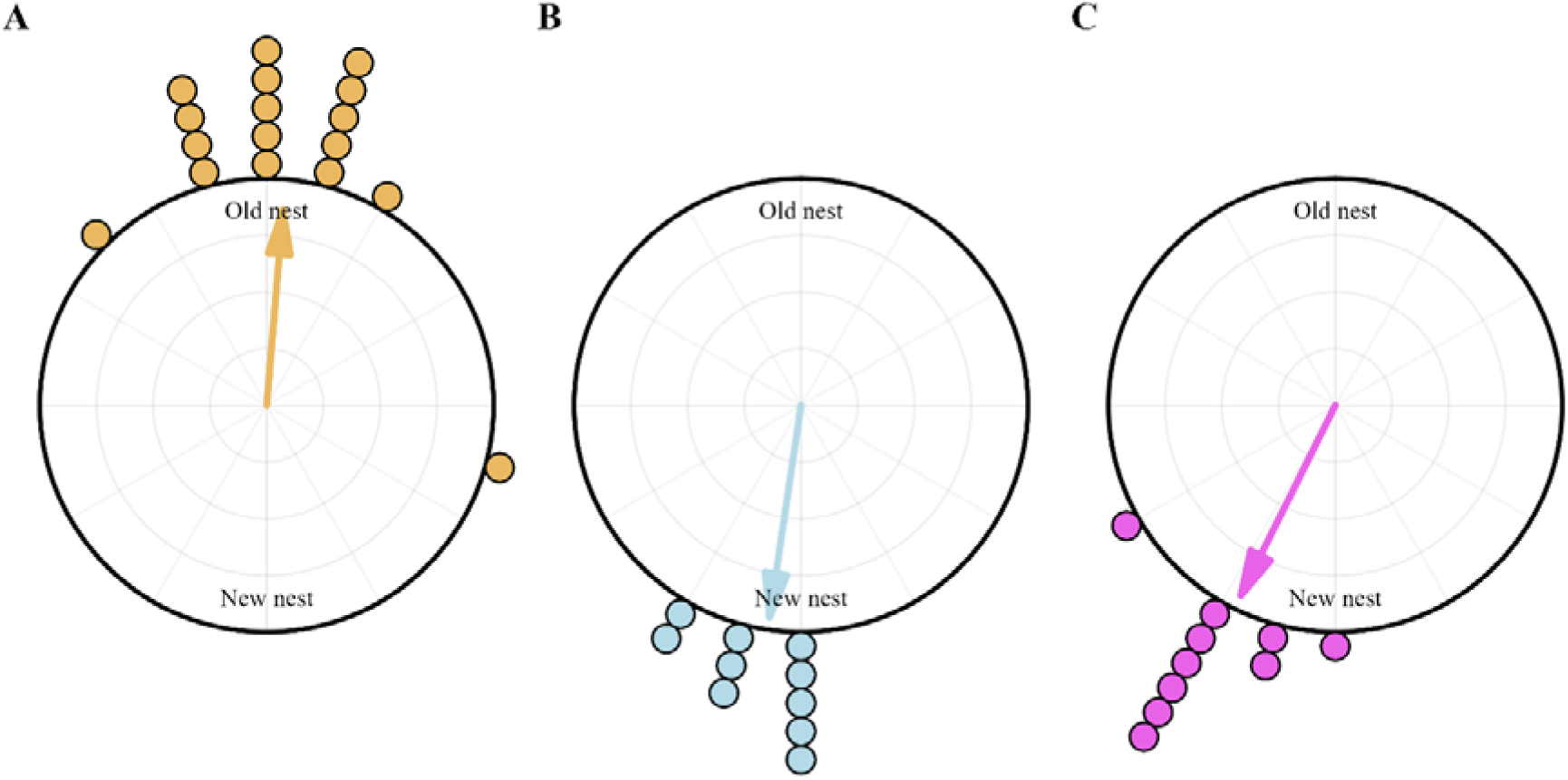
Circular histograms show the centroid direction of the ants during the exploration and transportation walks toward the Old nest (N ≥ 10) (A), toward the New nest (N = 10) (B) and Transportation (N = 10) (C) walks. In the histograms, the goal direction is set at 0°. The arrows denote the length and direction of the mean vector.

**Figure 3.**
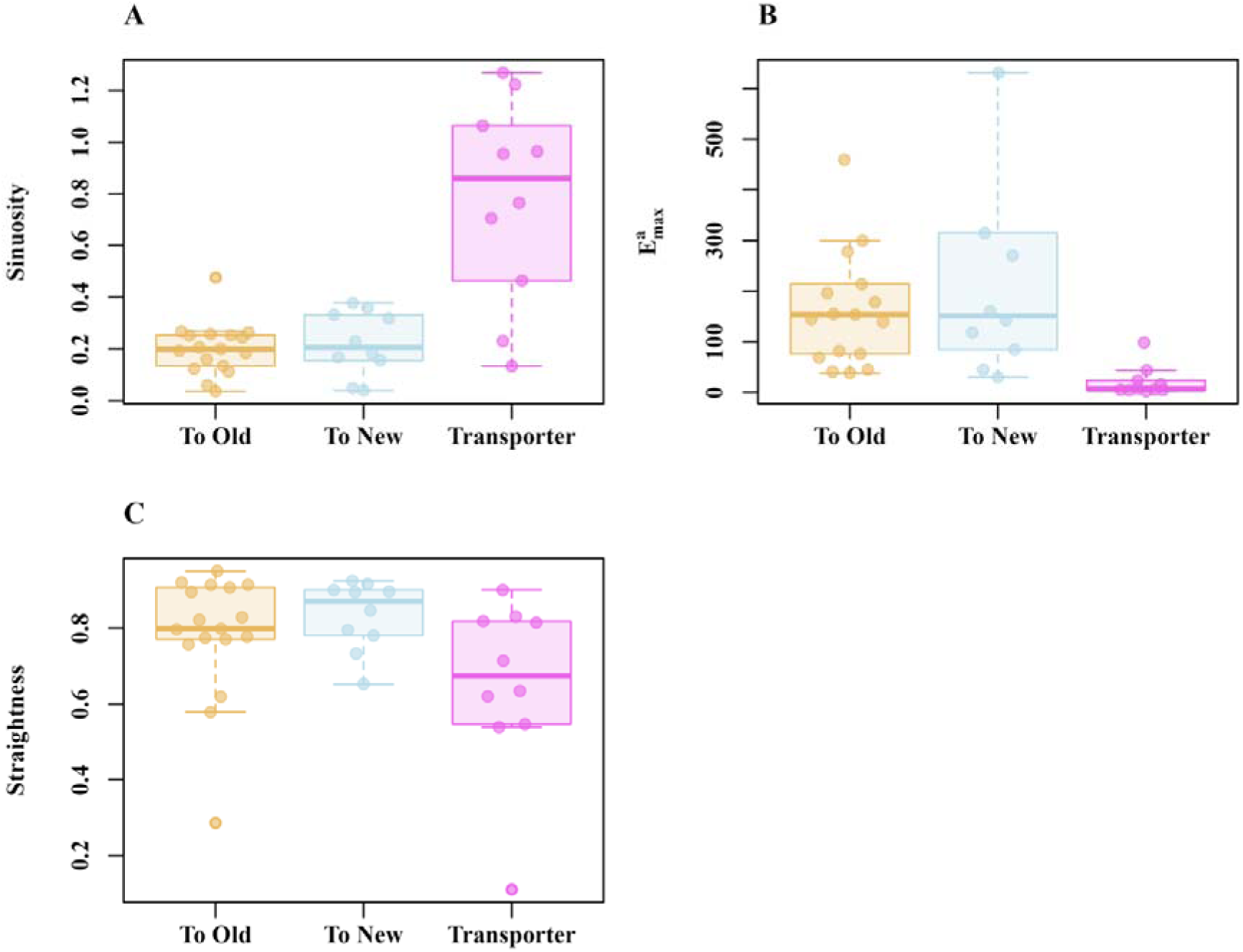
Path characteristics of ants carrying out different types of travel during the exploration and transportation walks toward the Old nest (N ≥ 10), New nest (N = 10) and Transportation (N = 10) walks: A) Sinuosity, B) E_max_ and C) straightness. Box plots display the median (line inside the box), interquartile range (box), and extreme values excluding outliers (whiskers). Individual data points are shown as dots. The data points falling more than 1.5 times the interquartile range beyond the upper or lower quartile are considered as outliers.

In path characteristics, we found more tortuosity (*sinuosity*) in Transportation walks than in either type of Exploration walks (Walk type: F _(2, 36)_ = 3.12, *p* ≤ 0.003; compared with New-to-Old Exploration walks: F _(1, 34)_ = 5.05, *p* ≤ 0.0001; compared with Old-to-New Exploration walks: F _(1, 34)_ = 6.5, *p* ≤ 0.0001; Figure 3A). We also found that Transporters had more constrained movement in their step displacement (*E*^a^_max_) (Walk type: F _(1, 36)_ = 3.74, *p* ≤ 0.0005; New nest vs. Transporter: F _(1, 34)_ = 36.1, *p* = 0.005; and Old nest vs. Transporter: F _(1, 34)_ = 32.8, *p* = 0.005) (Figure 3B). Transporters showed lower straightness numerically (Figure 3C). Although there is a suggestion of a trend, we did not find any significant differences between groups for our *straightness* measure (F _(2, 36)_ = 1.9, *p* = 0.057). In summary, the overall path analysis results indicate that transporters had more pronounced twists and turns in their path characteristics compared to ants executing exploratory walks.

When transporting resources to the new nest, ants walked more slowly than during either type of exploration walk (Figure 4). Typically, transporters carry larvae, pupae, callow, and some naive ants in their mandibles to the new nest. In 8 out of 10 observations of ants carrying brood, transporters were observed lifting the brood in their mandibles without touching the ground in their entire trip to the new nest. In the other 2 observations, transporters were noticed to touch the ground only occasionally. The slower speed is consistent with their higher sinuosity measure, as transporters exhibited more moment-to-moment turns that were accompanied by reduced speed during the walks compared to both types of exploration walks (Figures 4A–4D, blue lines showing the orientations during the walk). We found significantly lower speed in Transporters (Walk type: F _(2, 36)_ = 5.13, *p* ≤ 0.0005) than Exploration walks (Figure 4E). The post-hoc comparisons showed that Transporters moved more slowly compared to ants performing both Exploratory walks (Old nest vs. Transporter (F _(1, 34)_ = 2.9, *p* ≤ 0.001) and New nest vs. Transporter (F _(1, 34)_ = 2.5, *p* ≤ 0.001).

**Figure 4.**
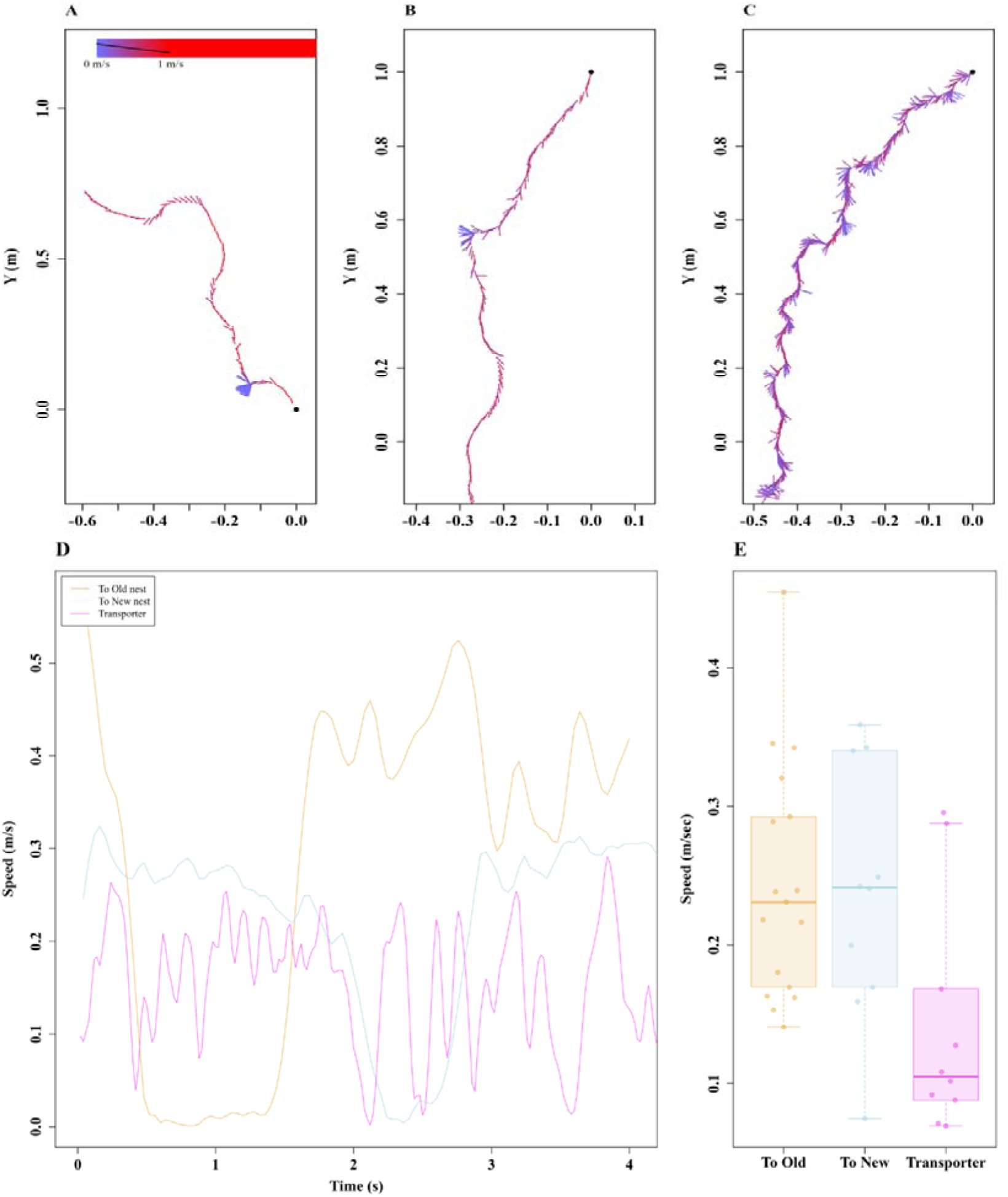
Speed variation in walks during different types of travel. The quiver plots display the speed variation during the walk. Line segment directions indicate direction of travel at each recorded point in time. Longer line segments indicate that the ant spent a longer time travelling in the indicated direction than short line segments (i.e. was moving more slowly). Segment color indicates speed. The three examples represented are: a walk to the Old nest (A), a walk to the New nest (B), and a walk transporting resources (C). Three examples of the speed of single ants plotted against time for the three types of travel (A, B, C). The box plots show speed values averaged across the recorded path in each type of travel (E). The boxes indicate the median and quartiles, while the whiskers show extreme values excluding outliers. Data points falling more than 1.5 times the interquartile range beyond the upper or lower quartile (none in the panel) are considered as outliers.

### Relearning behavior

During the relocation, we observed walks of foragers at both the Old and New nests that are similar to learning walks. Here we compared the relearning walks with actual first learning walks of the same species. During the relearning walks, ants stayed near the nest, walked in paths with many twists and turns to the left and right and ended back at the nest, which resembles the learning walks (Figures 5A, 5B and 5C). During the Old-nest relearning walks, ants leaving the nest travelled within a 15-cm distance mostly in the general direction of the New nest. Every single ant’s centroid on the walk lay consistently in a direction toward the New nest (S. Figure 2). Differences in path characteristics are found across these three types of walks. Firstly, during the Old-nest relearning walks, ants walked slower (Figure 5D); as a result, the GLM model found significant differences across types of walks (Walk type: F _(2, 27)_ = 13.15, *p* ≤ 0.001). The post-hoc comparisons showed significantly higher speeds at the New nest than at the Old nest (F _(1, 29)_ = 3.72, *p* ≤ 0.002), but no significant differences between relearning walks at the New nest and learning walks (F _(1, 29)_ =1.05, *p* = 0.58) and between relearning walks at the Old nest and learning walks (F _(1, 29)_ = 2.72, *p* = 0.02). Other walk characteristics also varied across walk types (Figure 6). The total area around the nest covered by the ants from the start of the trip to the end of the trip was significantly different among the walks (Walk type: F _(2, 27)_ = 10.54, *p* = 0.005; Figure 6A). The post-hoc comparisons showed a significantly smaller area on relearning walks at the Old nest than on learning walks (F _(1, 29)_ = 3.3, *p* ≤ 0.007), but no significant differences between relearning walks at the New nest and learning walks (F _(1, 29)_ = 1.26, *p* = 0.07) or between relearning walks at New and Old nests (F _(1, 29)_ =1.04, *p* = 0.55). The distance of the farthest point of the walk from the nest entrance also differed significantly across types of walks (Walk type: F _(2, 27)_ = 28.96, *p* ≤ 0.005; Figure 6B). The post-hoc comparisons showed significantly longer maximum distances at the New than at the Old nest (F _(1, 29)_ = 5.02, *p* ≤ 0.0001), and longer maximum distances on learning walks than on relearning walks at the Old nest (F _(1, 29)_ = 4.07, *p* ≤ 0.001), but no significant difference between relearning walks at the New nest and learning walks (F _(1, 29)_ =1.002, *p* = 0.57). Even though ants covered less area and did not move as far from the Old nest, they spent a longer duration performing relearning walks compared to ants at the New nest (Figure 6C). The post-hoc comparisons showed significantly longer walk durations on relearning walks at the Old nest than at the New nest (F _(1, 29)_ = 3.12, *p* ≤ 0.001), and longer durations on relearning walks at the New nest than on learning walks (F _(1, 29)_ = 2.56, *p* ≤ 0.001), but no significant differences between relearning walks at the Old nest and learning walks (F _(1, 29)_ =1.022, *p* = 0.47).

**Figure 5.**
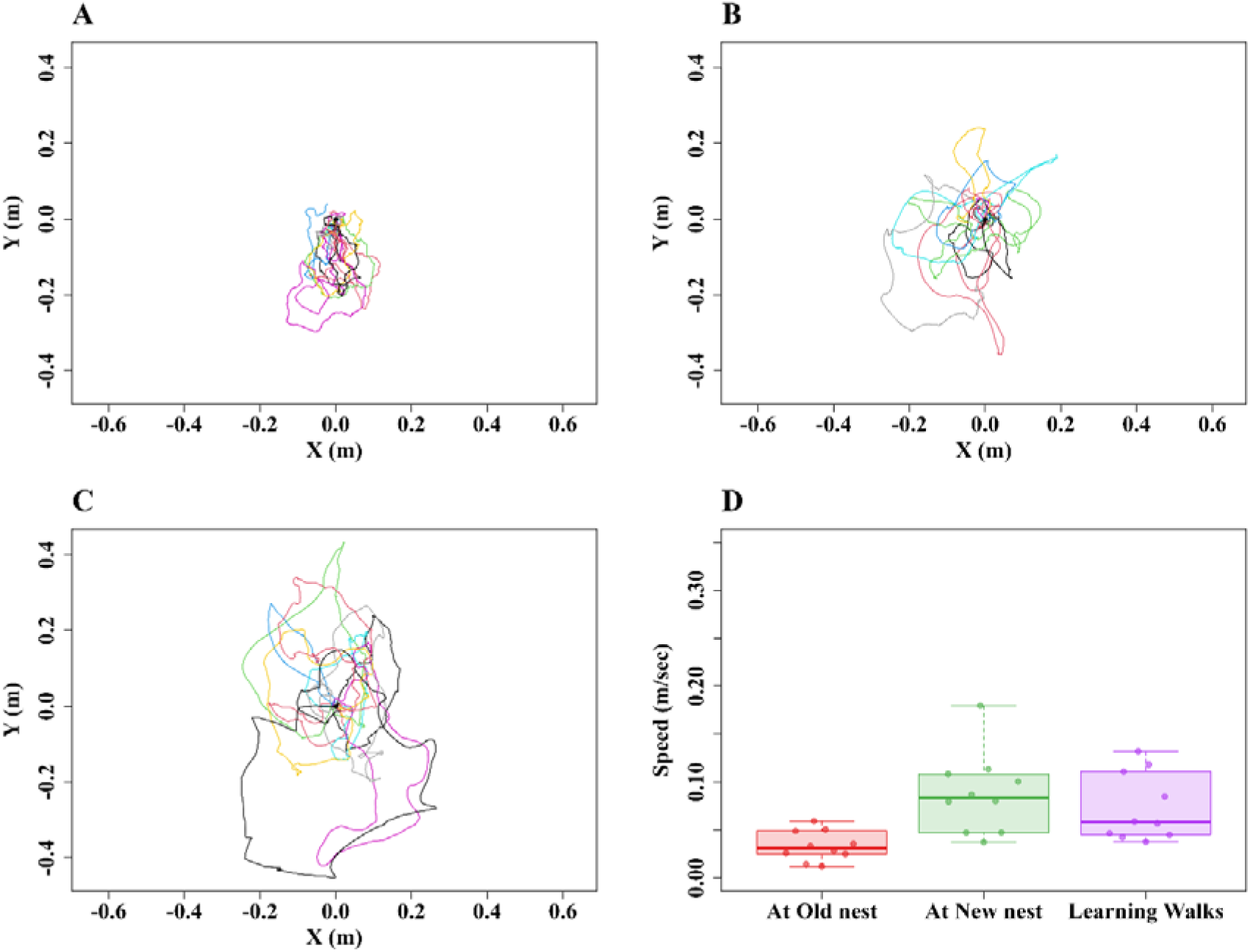
Relearning-walk and learning-walk trajectories and speed characteristics. The line graphs show the relearning walks at A) the Old nest, B) the New nest, and C) learning walks. The nest is located at (0, 0). D) Box plots showing the distribution of speed in each observed activity. The boxes indicate the median and quartiles, while the whiskers show extreme values excluding outliers. Data points falling more than 1.5 times the interquartile range beyond the upper or lower quartile (none in the panel) are considered as outliers.

**Figure 6.**
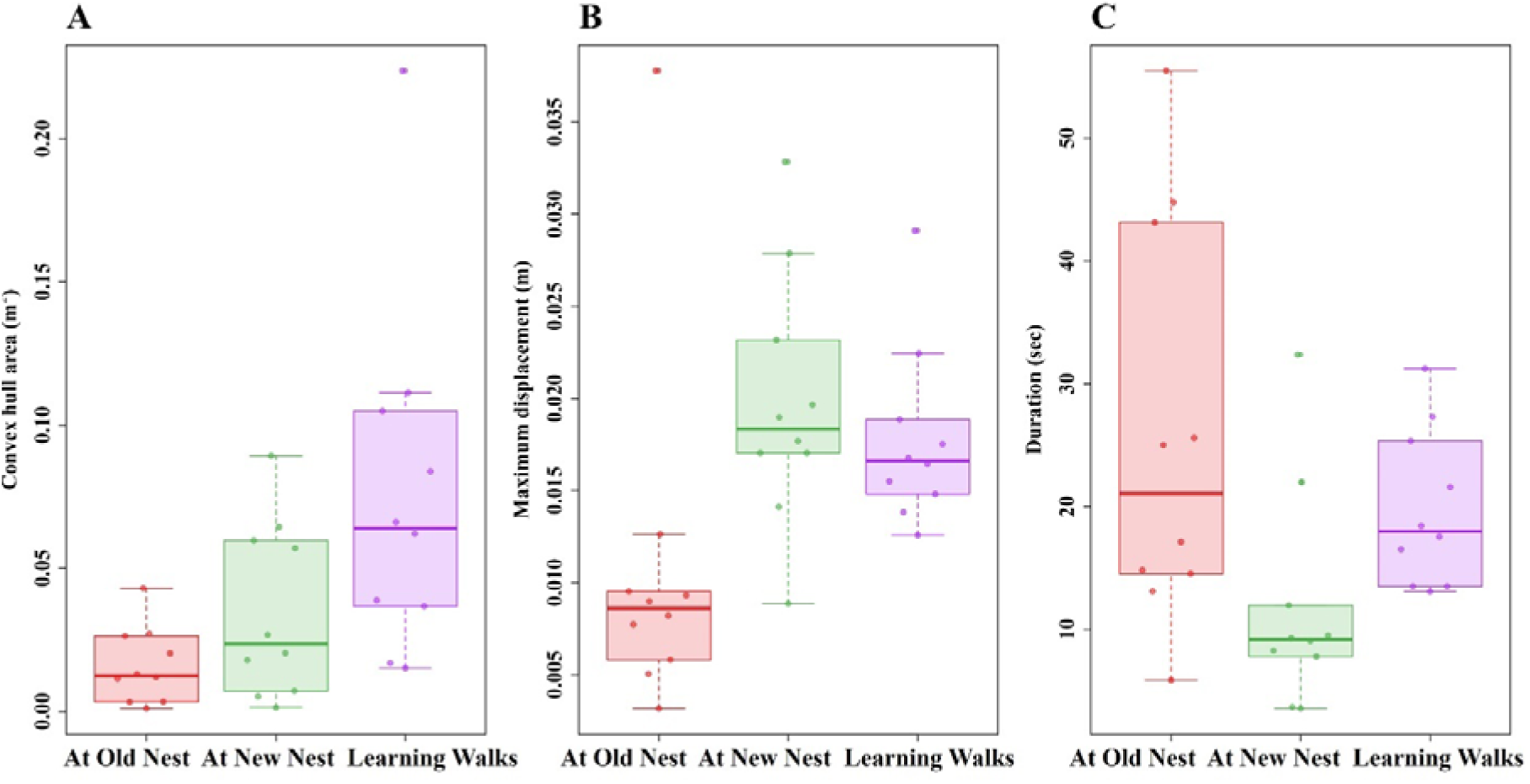
Characteristics of relearning walks at the Old and New nests and of learning walks. A) Convex hull area, B) maximum displacement from the nest, C) duration of the walk. The boxes indicate the median and quartiles, while the whiskers show extreme values excluding outliers. Each point represents a single trajectory. The data points falling more than 1.5 times the interquartile range beyond the upper or lower quartile are considered as outliers.

## Discussion

The phenomenon of nest relocation in solitarily foraging desert-ant colonies, as well as their behavioral response to nest damage caused by heavy rainfall, has not been previously investigated in any field-based or laboratory-based experiments. In the current field-based observations of the natural nest-relocation behavior of *M. bagoti,* we identified five types of behaviors: old-nest relearning walks, then old-to-new- and new-to-old-nest exploration, followed by old-to-new-nest transportation, then finally new-nest relearning walks. Notably, relearning at the Old nest showed directed movement, while relearning at the New nest exhibited a more uniform directional distribution. During transportation, in which ants were carrying brood or other colony resources, ants moved more slowly than during exploration.

In the walks between the two nests, while the exploratory walks between nests likely aided the learning of the route, the slower transportation walks appeared to efficiently transport resources. Transporters walked slower, with a higher degree of moment-to-moment curviness or sinuosity. Two non-mutually-exclusive reasons for these differences are that the resources being transported are more valuable to the colony than the pieces of food carried by foragers, and the ants are impeded by carrying materials in their mandibles. The slower speed might reflect the ants strategically reducing the chances of dropping or damaging the goods. We currently do not know what decisional and neurobiological processes might underlie this idea of taking care in walking. Perhaps a more likely cause is the material that an ant is carrying on transportation walks impedes travel. From other as yet unpublished observations on these ants, we do not think that it is the weight that causes ants to slow down. The material that an ant is transporting, however, is held in front of the ant and changes the panoramic view, especially in front of the travelling ant, and view changes cause changes in gait (unpublished observations). The side-to-side oscillations exhibited by transporters may be an attempt to view the scene in front unobstructed. Further tests are needed to examine these various interpretations.

When it comes to the relearning walks, those at the Old nest might be important for later transportation of resources to the new nest. The sequence of Old-nest relearning walks followed by exploration between nests supports such an interpretation. So is the fact that relearning walks at the Old nest are concentrated in the general direction of the New nest (Figure 4, S. Figure 2). New-nest relearning behaviors, on the other hand, suggest that their purpose is to enable future homing, such as after foraging. Walks at the new nest had a uniform directional distribution around the nest, covering all directions around the nest. Both Old- and New-nest relearning walks resembled the learning walks of naive ants to an extent, but differences in walk characteristics were found. The relearning walks at the New nest were faster, covered a larger area, and were not directed toward the Old nest. Relearning walks at the New nest resembled the first learning walks of future *M. bagoti* foragers, covering a similar area and maximum displacement. Future foragers gradually learn the foraging direction by increasing the area covered on learning walks with successive trips (Deeti & Cheng, 2021a; Deeti et al., 2024b). This strategy is also observed in the tree-foraging bull ant *Myrmecia crosslandi* (Jayatilaka et al., 2018). At the Old nest, relearning walks contributed to the development of the one route leading to the New nest, while at the New nest, ants were attempting to learn the surrounding scene to return to their new home from various directions. The two types of relearning walks were geared at successful nest relocation at the Old nest and spatial knowledge acquisition at the New nest. These different functions likely explain the differences in characteristics between relearning walks at the Old and New nests.

While exploratory or scouting behavior for finding a suitable nest site and optimal food resources is well known in insects (Doran & Larsen, 2016; Janson et al., 2008), how these animals coordinate the transportation of brood and other colony resources to the new nest site was largely unknown. We have shown that workers explore the route between the old and new nests before transporting resources to the new nest. Since desert ants do not rely on pheromone trails for navigation (Graham & Cheng, 2009; Wehner et al., 2014; Wystrach et al., 2014), these ants may be familiarizing themselves with the route between the old and new nest. This familiarization could develop their route memory and learning of the nearby environmental cues. Alternatively, these exploratory walks may help ants identify potential hazards and predator risks in the new environment, which could otherwise interfere with relocation and nest establishment. Thirdly, it is possible that the ants are compensating for the large distance between the old and new nests by engaging in these exploration walks before transporting resources. Foraging distances are often shorter than the 20-m distance between nests that we observed (Muser et al., 2005). These possible explanations are not mutually exclusive, and further research on this topic is needed to evaluate them.

Our study shows that learning and navigation play key roles in nest relocation in this species (Deeti & Cheng, 2021b; Schultheiss et al., 2010b). In one earlier study, a *M. bagoti* worker was observed dragging her abdomen across the sandy ground to direct the colony members toward the new nest (Schultheiss et al., 2010). However, this study did not definitively establish whether all exploratory or transporting ants lay intermittent odor trails. The desert climate destroys pheromones quickly, so that even with odor trails as guides, ants might well need to learn visual cues in addition to travel the route regularly. Our study shows these desert ants doing just that. Future studies should examine how many relearning walks are required before starting the relocation.

Nest relocation is uncommon for many solitarily foraging desert ant colonies with a large workforce, such as the Australian desert ant *Melophorus bagoti*, which, based on the personal observations, typically does not move nests. For any ant colony, however, various exogenous and endogenous factors may cause them to relocate. In the current study, heavy rainfall triggered the relocation of the observed desert ant colony to a new site 20 m away from their old nest. While two previous studies have reported instances of nest relocation in this species (Schultheiss & Cheng, 2010; Deeti & Cheng, 2021a), nest disturbance from field experimentation, such as setting up feeders and partitioning areas around nests, usually does not lead to nest movement. Heavy rainfall has also been associated with relocation in nocturnal bull ants *Myrmecia midas* colonies (Deeti et al., 2024a). Record rains in NSW in 2022 led to deterioration of nest chambers in a sizeable minority of nests at the field site. Even though some social insects like termite colonies are not sessile and shift locations even in the absence of significant changes in environmental conditions (McGlynn, 2012), the current nest relocation in the red honey ants is likely to have been triggered by structural damage to the nest chambers caused by the heavy rain.

One limitation of the current study is that it is based on a single nest, unlike the multiple relocation events observed in bull ants (Deeti et al., 2024a). Given the evidence that flooding can cause nest relocation, it is possible and legal to experimentally flood nests and then observe nest relocation. Although legal, and although the ants are common in Central Australia, we would still find it unethical to deliberately cause the destruction of a red-honey-ant nest.

In sum, this study investigated the nest relocation behavior and response to nest damage in the solitarily foraging desert ant *Melophorus bagoti,* particularly focusing on their behavioral patterns and learning processes during relocation. Heavy rainfall causing structural damage to nest chambers likely triggered the observed nest relocation. The field-based observations revealed a sequence of behaviors, from old-nest relearning walks to exploration between old and new nests to transportation walks ferrying resources to the new nest to new-nest relearning walks. Notably, transporters exhibited slower and more sinuous movements when carrying brood or resources, suggesting a careful approach in transportation. The relearning walks at the old nest appeared to facilitate later resource transportation to the new nest, while relearning walks at the new nest seemed geared toward spatial knowledge acquisition. The study highlights the importance of exploratory walks in familiarizing ants with the route between old and new nests, contributing to route memory and identification of environmental cues. The lack of reliance on pheromone trails in desert ants highlights the significance of visual cues and regular route travel for successful relocation. The findings emphasize the role of learning and navigation in nest relocation. Future studies should explore the number of relearning walks required before initiating relocation. Overall, this study revealed some of the complex behavioral processes involved in nest relocation in *Melophorus bagoti,* offering insights into the ant’s adaptive responses to environmental changes and challenges.

## Acknowledgements

We acknowledge the traditional custodians of the land upon which this research was conducted, the Arrernte people. Their culture and customs have nurtured and sustained this land since the Dreamtime and continue to do so today. We pay our respects to their Elders past and present. We thank the Centre for Appropriate Technology at Alice Springs, Australia for letting us work on their property and providing some storage space, and the CSIRO Arid Zone Research at Alice Springs for administrative support. We are also thankful to Cody Freas for helping me during the field trip.

## Author contributions

Experimental design: SD. Data collection: SD. Data analysis: SD and DJM. Writing: SD, TM and KC.

## Ethics standards

Australia has no ethical regulations regarding work with insects. The study was non-invasive and no long-term aversive effects were found on the nests or on the individuals studied.

## Data availability

Supplementary videos, Excel file and R scripts are available at Open Science framework: and DRYAD: https://doi.org/10.5061/dryad.g1jwstqz5.

---

**S. Figure 1.**
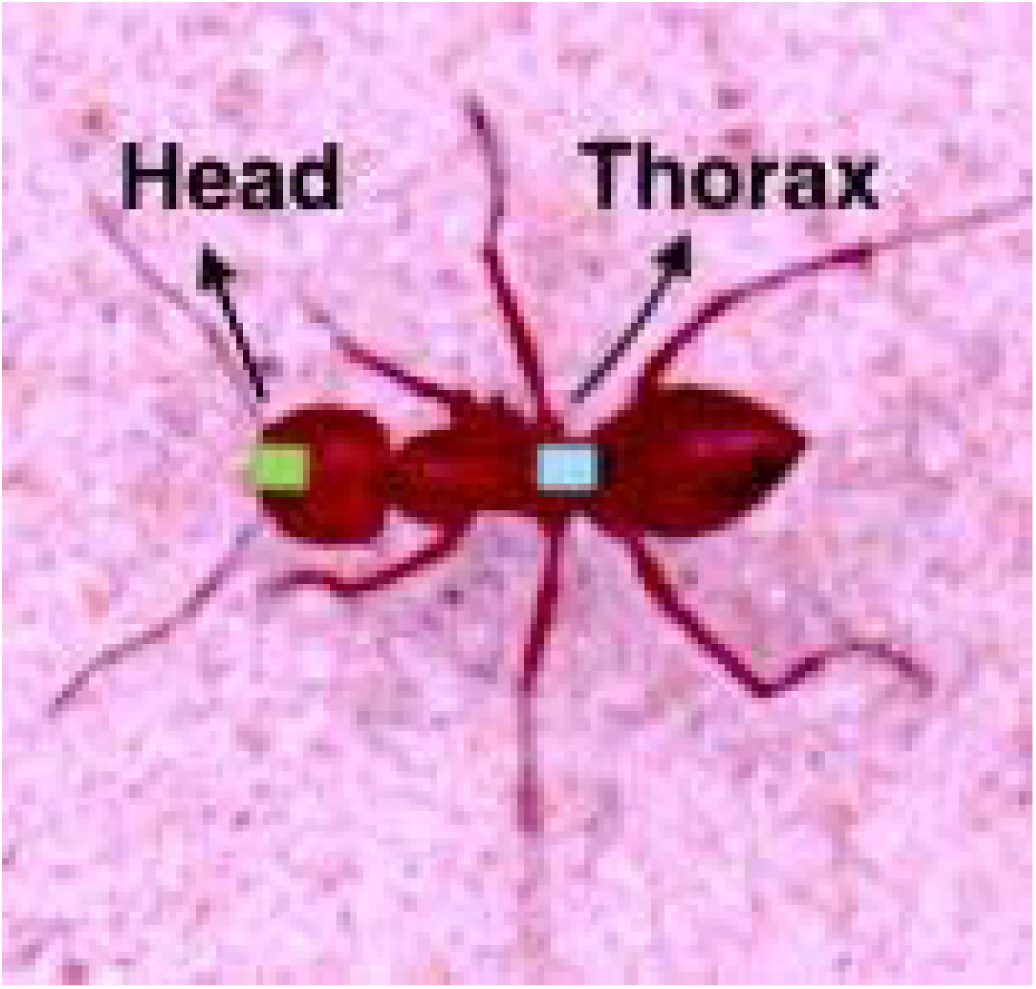
The body positions used to annotate ants on video records: Front of the head and Mid thorax.

**S. Figure 2.**
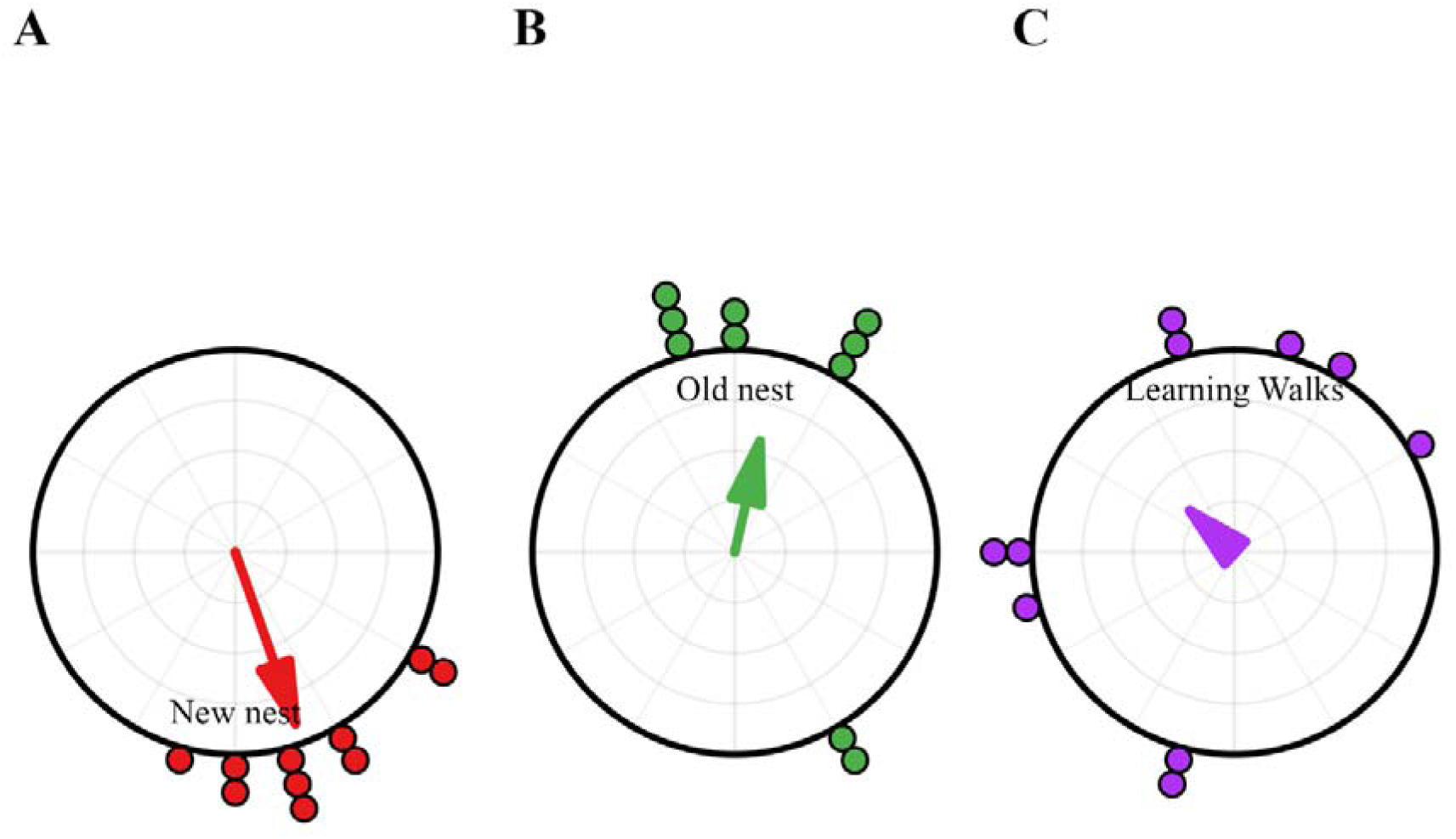
Circular histograms show the centroid direction of the ants during the relearning walk at Old nest (A), relearning walks at the New nest (B) and naïve learning walks. In the histograms, the nest direction is set at 0°. The arrows denote the length and direction of the mean vector. The Rayleigh uniformity test and the distribution of headings with the *V* test showed significance in centroid heading toward the New nest at Old nest relearning walks (Z = 9.16, *p* ≤ 0.001; *V* test: Z = 4.72, *p* ≤ 0.001), whereas the New nest relearning walks (Z = 1.16, *p* = 0. 076; *V* test: Z = 1.02, *p* = 0.81) and learning walks (Z = 0.16, *p* = 0.41; *V* test: Z = 0.74, *p* = 0.24) showed random distribution in their centroid direction.

